# Macroscale structural changes of thylakoid architecture during high light acclimation in *Chlamydomonas reinhardtii*

**DOI:** 10.1101/2021.06.26.450046

**Authors:** Mimi Broderson, Krishna K. Niyogi, Masakazu Iwai

## Abstract

Photoprotection mechanisms are ubiquitous among photosynthetic organisms. The photoprotection capacity of the green alga *Chlamydomonas reinhardtii* is correlated with protein levels of stress-related light-harvesting complex (LHCSR) proteins, which are strongly induced by high light (HL). However, the dynamic response of overall thylakoid structure during acclimation to growth in HL has not been characterized. Here, we combined live-cell super-resolution microscopy and analytical membrane subfractionation to investigate macroscale structural changes of thylakoid membranes during HL acclimation in *C. reinhardtii*. Subdiffraction-resolution bioimaging revealed that overall thylakoid structures became thinned and shrunken during HL acclimation. The stromal space around the pyrenoid also became enlarged. Analytical density-dependent membrane fractionation indicated that the structural changes were partly a consequence of membrane unstacking. The analysis of both an LHCSR loss-of-function mutant*, npq4 lhcsr1*, and a regulatory mutant that over-expresses LHCSR, *spa1-1*, showed that structural changes occurred independently of LHCSR protein levels, demonstrating that LHCSR was neither necessary nor sufficient to induce the thylakoid structural changes associated with HL acclimation. In contrast, *stt7-9*, a mutant lacking a kinase of major light-harvesting antenna proteins, had a distinct thylakoid structural response during HL acclimation relative to all other lines tested. Thus, while LHCSR and the antenna protein phosphorylation are core features of HL acclimation, it appears that only the latter acts as a determinant for thylakoid structural rearrangements. These results indicate that two independent mechanisms occur simultaneously to cope with HL conditions. Possible scenarios for HL-induced thylakoid structural changes are discussed.

## INTRODUCTION

Understanding natural photosynthetic systems is crucial for developing efficient technologies for bioenergy. In green algae and plants, photosynthetic electron transport is initiated by the absorption of light energy mainly by light-harvesting complex (LHC) proteins in chloroplast thylakoid membranes (Jansson, 1999, Wobbe *et al.*, 2016). Light energy is essential for photosynthesis, but an excess amount of absorbed energy can cause photooxidative damage, which depresses photosynthetic efficiency and growth (Aro *et al.*, 1993, Li *et al.*, 2009, Pinnola and Bassi, 2018). Therefore, photosynthetic organisms have evolved photoprotective mechanisms. One of the most important mechanisms is nonphotochemical quenching (NPQ) of chlorophyll (Chl) fluorescence, which harmlessly dissipates excess excitation energy as heat (Muller *et al.*, 2001, Ruban, 2016). The fast-acting component of NPQ, qE, is tightly regulated by a proton gradient across the thylakoid membrane (ΔpH). The ΔpH is determined by the rate of water oxidation catalyzed by the oxygen-evolving complex of photosystem II (PSII) and the Q-cycle of the cytochrome *b*_6_*f* complex in the light (Kramer *et al.*, 2004). This light-induced increase in ΔpH is essential for not only generation of ATP by ATP synthase but also the feedback induction of qE to minimize photooxidative damage in thylakoid membranes under fluctuating light conditions in natural environments.

An increased flow of photosynthetic electrons from PSII also causes the accumulation of reduced plastoquinone (PQ) pool in thylakoid membranes, which triggers another component of NPQ, qT or state transitions. The reduced PQ pool activates the Stt7/STN7 kinase, which phosphorylates LHC proteins of PSII (LHCII) to induce redistribution of excitation energy from PSII to photosystem I (PSI) via increased connectivity between LHCII and PSI (Minagawa, 2011, Rochaix, 2011, Goldschmidt-Clermont and Bassi, 2015). The redistribution of excitation energy between the two photosystems prevents excessive excitation of PSII, while PSI converts excitation energy more efficiently (Trissl and Wilhelm, 1993). The time scale of activation of state transitions is relatively longer than qE, because it involves dynamic protein reorganization within thylakoid membranes.

NPQ is ubiquitous among oxygenic photosynthetic organisms, although the molecular processes are different (Niyogi and Truong, 2013, Pinnola, 2019). In the model unicellular green alga *Chlamydomonas reinhardtii*, the stress-related LHC proteins, called LHCSRs, are essential for the qE component of NPQ (Peers *et al.*, 2009). Similar to components of the major LHC antenna system, LHCSR is a membrane protein with three transmembrane helices and contains Chls and carotenoids (Bonente *et al.*, 2011, Liguori *et al.*, 2013). The qE facilitated by LHCSR is regulated by the protonation state of acidic residues on the lumenal side of the protein, and this protonation state responds to changes in ΔpH under high light (HL) conditions (Ballottari *et al.*, 2016, Dinc *et al.*, 2016, Troiano *et al.*, 2021). Intriguingly, LHCSR protein level in thylakoid membranes is very limited under low light (LL) conditions (Petroutsos *et al.*, 2011, Maruyama *et al.*, 2014). Recently, it has been shown that the expression of NPQ-related genes, including *LHCSRs*, is negatively regulated by a conserved E3 ubiquitin ligase complex, which targets the transcriptional activators for qE gene expression in LL for proteolysis (Aihara *et al.*, 2019, Gabilly *et al.*, 2019, Tokutsu *et al.*, 2019). Under HL conditions, the E3 ubiquitin ligase activity is inactivated by a photoreceptor-dependent mechanism, which allows the transcriptional activation of the qE-related genes. The inhibition of the E3 ubiquitin ligase activity by a mutation in its component SPA1 results in the accumulation of LHCSR proteins even under LL conditions (Gabilly *et al.*, 2019, Tokutsu *et al.*, 2019). Thus, the induction of high NPQ activity in *C. reinhardtii* is tightly regulated during HL acclimation by both transcriptional and post-translational control of LHCSR activity in thylakoid membranes.

The molecular genetics and biochemistry of HL acclimation mechanisms in *C. reinhardtii* have been extensively studied (Allorent and Petroutsos, 2017, Rochaix and Bassi, 2019). Yet, the dynamics of the thylakoid membrane macrostructural response to HL acclimation have not been explored. In this study, we combined subdiffraction-resolution microscopy and analytical membrane subfractionation to investigate the macroscale structural changes of thylakoid architecture during HL acclimation in *C. reinhardtii*. We observed a drastic change in thylakoid structure during HL acclimation by using subdiffraction-resolution imaging analysis. We also observed changes in the densities of isolated thylakoid membranes, which were analytically separated by density-dependent fractionation. To investigate a correlation between the structural changes of thylakoid membranes and qE capacity, we used the *npq4 lhcsr1* mutant lacking LHCSR proteins (Ballottari *et al.*, 2016) and the *spa1-1* mutant (Gabilly *et al.*, 2019), in which LHCSR proteins are accumulated under LL. Regardless of the accumulation of LHCSR proteins, both mutants exhibited similar structural changes to wild-type (WT) after HL acclimation, demonstrating that the HL-induced changes in thylakoid structures occur independently of LHCSR function. On the other hand, the *stt7-9* mutant, which is deficient in state transitions, did not undergo the macroscale structural changes that occur in WT after HL acclimation. The two combined technical approaches revealed that the aspects of the macroscale membrane architecture and the membrane stacking can be characterized separately.

## RESULTS

To observe thylakoid membranes in *C. reinhardtii*, we used a confocal microscope with Airyscan, which is an array of 32 GaAsP photomultiplier tube detectors arranged in a hexagonal pattern and is capable of subdiffraction-resolution imaging through determining the microscope point spread function that is projected in the center of the array detectors (Huff *et al.*, 2017). The spatial resolution achievable by Airyscan microscopy is theoretically lower than that of structured illumination microscopy (SIM). However, it is reported that Airyscan microscopy performs better than SIM for samples with a lower signal-to-noise ratio (Sivaguru *et al.*, 2018). As compared to our previous study using SIM (Iwai *et al.*, 2018), Airyscan images contained no obvious artifact due to lower signal-to-noise ratios. Therefore, these two microscopes are complementary techniques to obtain subdiffraction resolution.

We first observed live *C. reinhardtii* cells acclimated to LL (~30 μmol photons m^−2^ s^−1^) using the Airyscan microscope. The Airyscan images showed typical thylakoid structures in the cup-shaped chloroplast very similar to the ones as previously observed using SIM (Iwai *et al.*, 2018). The LL-acclimated cells typically showed undisturbed thylakoid layers with relatively little space between the layers throughout the lobe regions (Figure 1a, b). By optical sectioning through the z-axis, it was possible to observe the membrane surface patterns (Figure 1c), which are usually difficult to obtain by electron microscopy (EM) observation. The Airyscan image revealed smooth fluorescence patterns indicating continuous membrane regions with occasional empty spaces. Layers of thylakoid membranes in the lobe and thylakoid tubules were also visible (Figure 1d, e). On the other hand, after HL (~350 μmol photons m^−2^ s^−1^) acclimation for 24 h, the overall thylakoid structures appeared to be thinner than that of the LL-acclimated cells. The thylakoid structures at the lobe showed fewer layers with more non-fluorescent spaces between the layers (Figure 2a-d). Similarly, the thylakoid structures at the base became thinned, and the space occupied by the pyrenoid became larger than that of the LL-acclimated cells. Optical sectioning showed that the surface pattern became rough and less dense due to the thinning of membranes (Figure 2e). Further, the lobe regions appeared to be shorter and sometimes partially disappeared.

**Figure 1.**
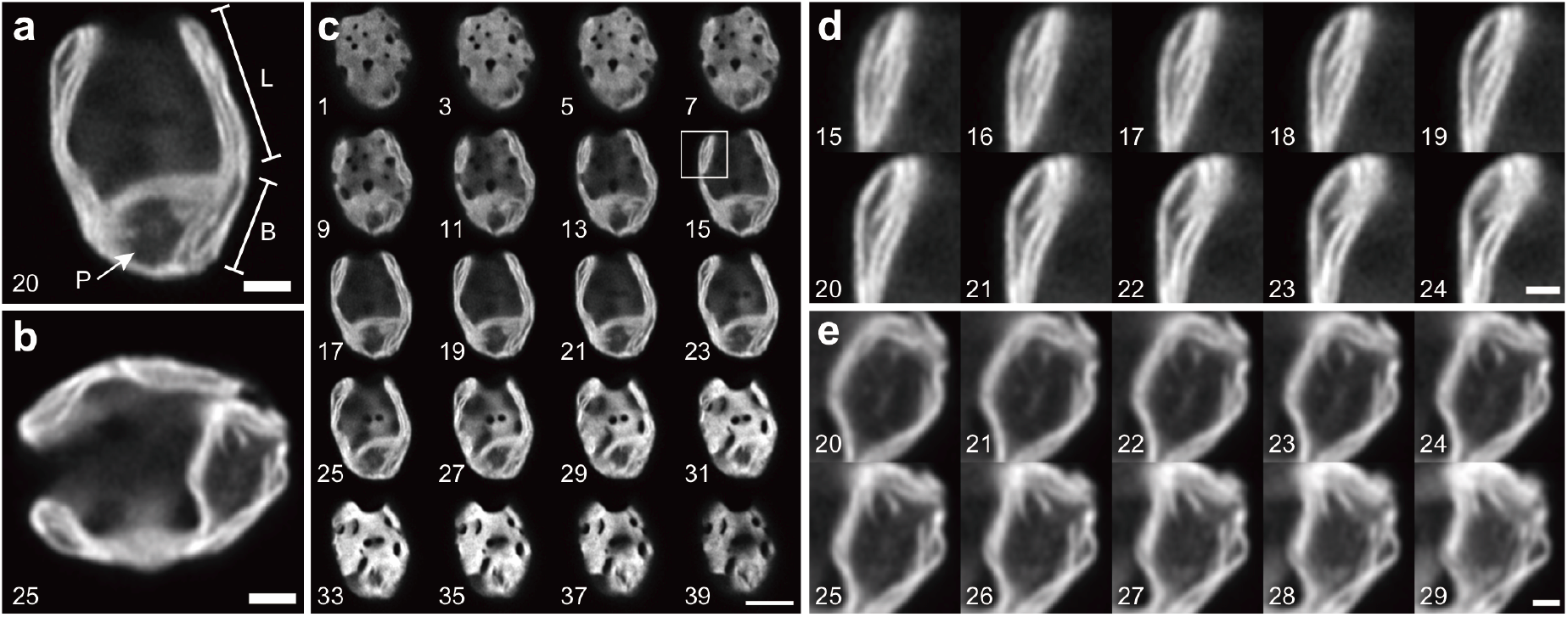
Airyscan images of *C. reinhardtii* acclimated to LL. **a, b.** Representative images of the LL-acclimated cells observed by Chl fluorescence. L, lobe; B, base; and P, pyrenoid. **c.** Optical serial sectioning of the cell (in **a**) through z-stack. **d.** Enlarged images for optical sectioning of a lobe region (a square in **c**), showing layers of thylakoid membranes. **e.** Enlarged images for optical sectioning of a pyrenoid region (in **b**), showing pyrenoid tubules (Engel *et al.*, 2015). Numbers indicate slice numbers of z-stack images. Scale bars = 2 μm (**a**, **b**), 5 μm (**c**), 1 μm (**d**, **e**).

**Figure 2.**
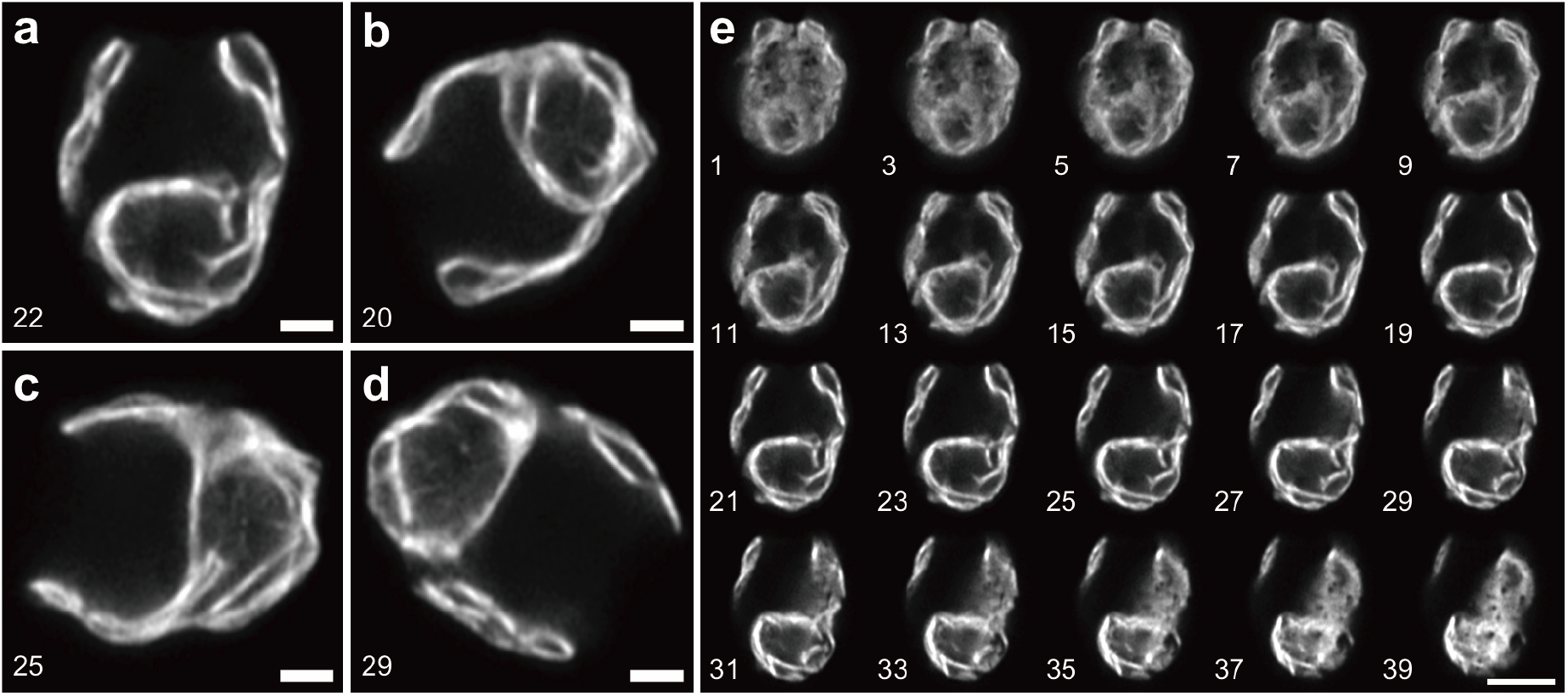
Airyscan images of *C. reinhardtii* acclimated to HL. **a-d.** Representative images of the HL-acclimated cells observed by Chl fluorescence. **e.** Optical serial sectioning of the cell (in **a**) through z-stack. Numbers indicate slice numbers of z-stack images. Scale bars = 2 μm (**a**-**d**), 5 μm (**e**).

Similar characteristics were observed under a transmission EM (TEM). The TEM images of the LL-acclimated cells revealed that thylakoid membranes were continuously undisturbed with narrow spaces between the membranes (Figure 3a). The membranes at the lobe regions were close to each other, and the ones at the base were also tightly associated near the pyrenoid (Figure 3b, c). The TEM images of the HL-acclimated cells clearly showed more stromal spaces between the membranes throughout the thylakoid structures (Figure 3d). In the lobe regions, there were larger spaces between the membranes, but also some membranes appeared to be associated tightly with each other (Figure 3e, g, h). Interestingly, the enlarged space for the pyrenoid is actually the stromal space around the pyrenoid, as the size of the pyrenoid did not significantly change under HL (Figure 3f). The membranes in the base regions also appeared more spaced out (Figure 3f, i). These TEM results demonstrated consistency with the Airyscan images of the thinner thylakoid structures and more space between membranes observed in the HL-acclimated cells.

**Figure 3.**
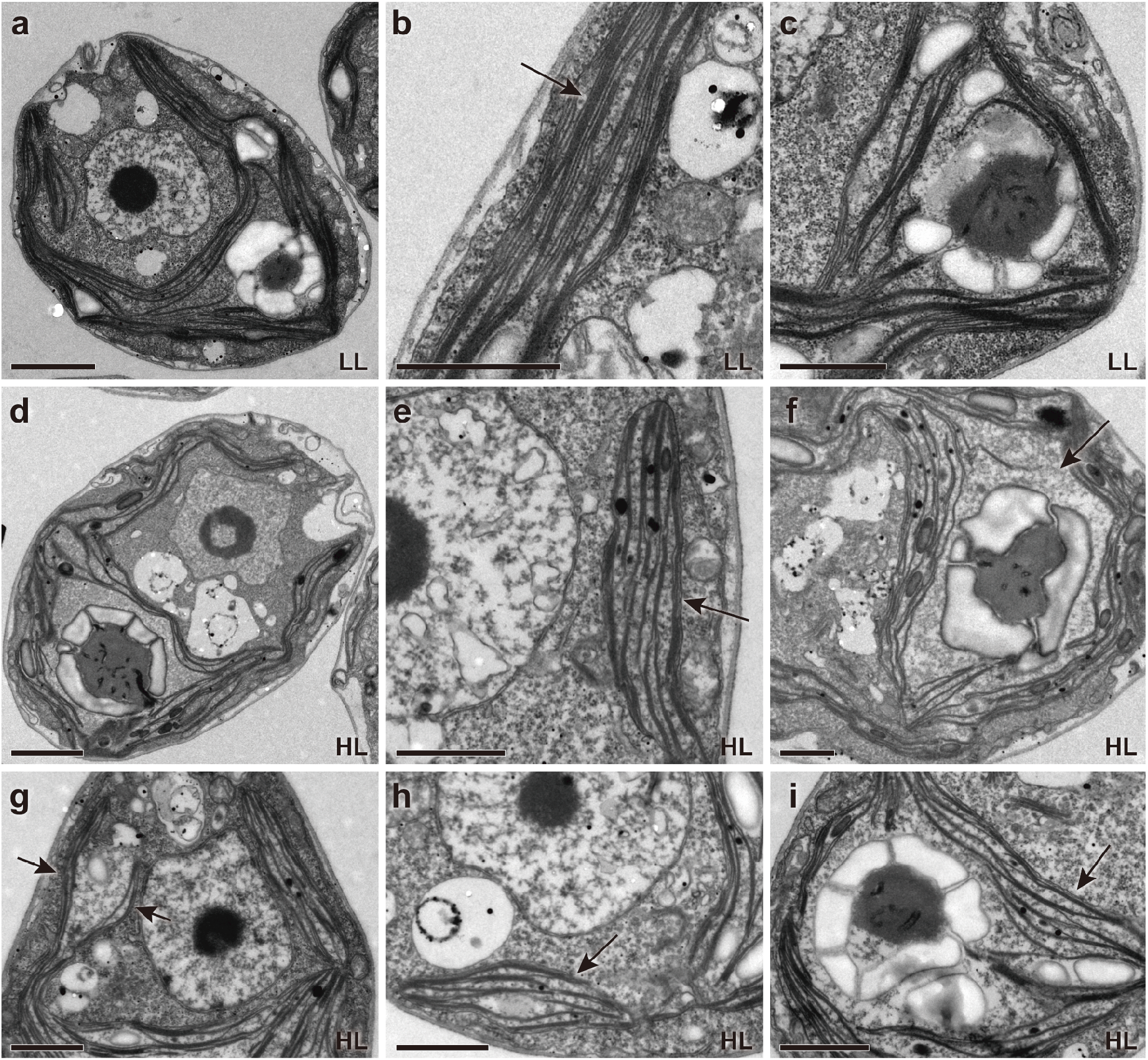
TEM images of *C. reinhardtii* acclimated to LL and HL. **a-c.** Representative images of the LL-acclimated cells, showing an overall cell structure (**a**), thylakoid membranes (indicated by an arrow) at a lobe region (**b**), and a base region (**c**). Representative images of the HL-acclimated cells, showing an overall cell structure (**d**), thylakoid membranes (indicated by an arrow) at lobe regions (**e, g, h**), and enlarged stromal spaces (an arrow in **f**) around the pyrenoid at base regions (**f, i**). Scale bars = 2 μm (**a**, **d**), 1 μm (**b, c, e-i**).

To gain more insight into the thylakoid membrane structures, we performed analytical density-dependent fractionation of isolated thylakoid membranes using sucrose step-wise gradient centrifugation. This experiment showed that the chloroplasts in the LL-acclimated cells contained a thick green band of the densest membrane fraction (T4) and a less distinct green band of a lighter membrane fraction (T3) (Figure 4a). In contrast, the chloroplasts in the HL-acclimated cells contained four green bands—the T4 fraction became least abundant, while the second lighter fraction (T2) and T3 fractions became abundant, and the lightest fraction (T1) became visible (Figure 4a). An increase of carotenoid was also visible above the T1 fraction. The increase in quantity of the lighter membrane fractions in the HL-acclimated cells is consistent with the imaging results that showed that overall thylakoid structures became thinned in the HL-acclimated cells (Figure 2). Surprisingly, an increase in T1-T3 fractions was already generated after 15 min of HL treatment (Figure 4b). A gradual increase in T1 and T2 was observed from 15 min to 1 h of HL treatment, while the accumulation of T4 declined. The membrane heights measured using atomic force microscopy (AFM) also indicated that the T4 fraction contained thicker membranes than T3, and the membranes in T1+T2 fractions were thinnest (Figure 4c). The changes in height reflects the number of membrane stacks, as thylakoid membranes in *C. reinhardtii* also form the appressed and non-appressed membrane regions. Recently, the lateral heterogeneity of the two photosystems has been shown by cryo-electron tomography; PSII is located in the appressed regions, while PSI is physically segregated into the non-appressed regions (Wietrzynski *et al.*, 2020). Chl fluorescence spectra at 77 K revealed that the T4 from LL contained the emission peaks for both PSII and PSI, but the T3 from HL showed much less PSI emission than the T1+T2 from HL (Figure 4d). These results suggest that the HL-induced thylakoid structural changes involve membrane unstacking.

HL induces both ΔpH and LHCSR expression, which are essential components for NPQ induction in *C. reinhardtii*. Given its central role in NPQ and high light acclimation, one hypothesized role of LHCSR has been the induction of thylakoid structural changes during HL acclimation. We tested this directly by using the *npq4 lhcsr1* mutant, which lacks LHCSR proteins and has low NPQ capacity even after HL acclimation (Ballottari *et al.*, 2016). Airyscan image analysis revealed that the structural changes of thylakoid membranes in *npq4 lhcsr1* were very similar to that of WT after HL acclimation—thinned lobe structures, more space between membranes, and a larger stromal space around the pyrenoid (Figure 5a-d). Analytical density-dependent fractionation of isolated thylakoid membranes also showed similar results— an increase in T1, T2, and T3 fractions and a decrease in T4 after HL acclimation (Figure 5e). These results indicate that the LHCSRs are not necessary for the thylakoid structural changes during HL acclimation. Additionally, the low NPQ capacity of HL-grown *npq4 lhcsr1* demonstrates that thylakoid membrane structural changes are not sufficient to induce qE in the absence of LHCSRs in *C. reinhardtii*.

**Figure 4.**
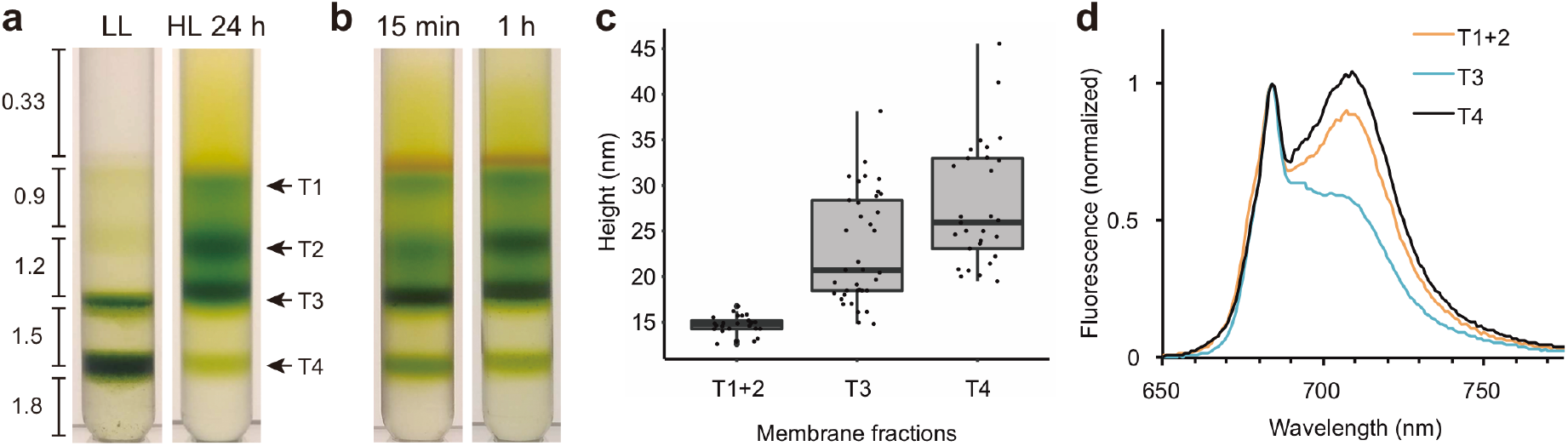
Analytical sucrose density-dependent membrane fractionation. **a.** The cells acclimated to LL and HL were gently disrupted (a total of 0.5 mg Chl) and separated by sucrose step-gradient centrifugation. Numbers indicate the molar concentration of sucrose (see Methods for details). Different membrane fractions were indicated as T1-T4. **b.** The results for the cells treated under HL for 15 min and 1 h. **c.** The height of isolated membranes in each fraction was measured by AFM. **d.** Chl fluorescence emission spectra at 77 K. Spectra were normalized at 684 nm. T1 and T2 (T1+2) and T3 were obtained from HL samples, and T4 was obtained from LL samples.

**Figure 5.**
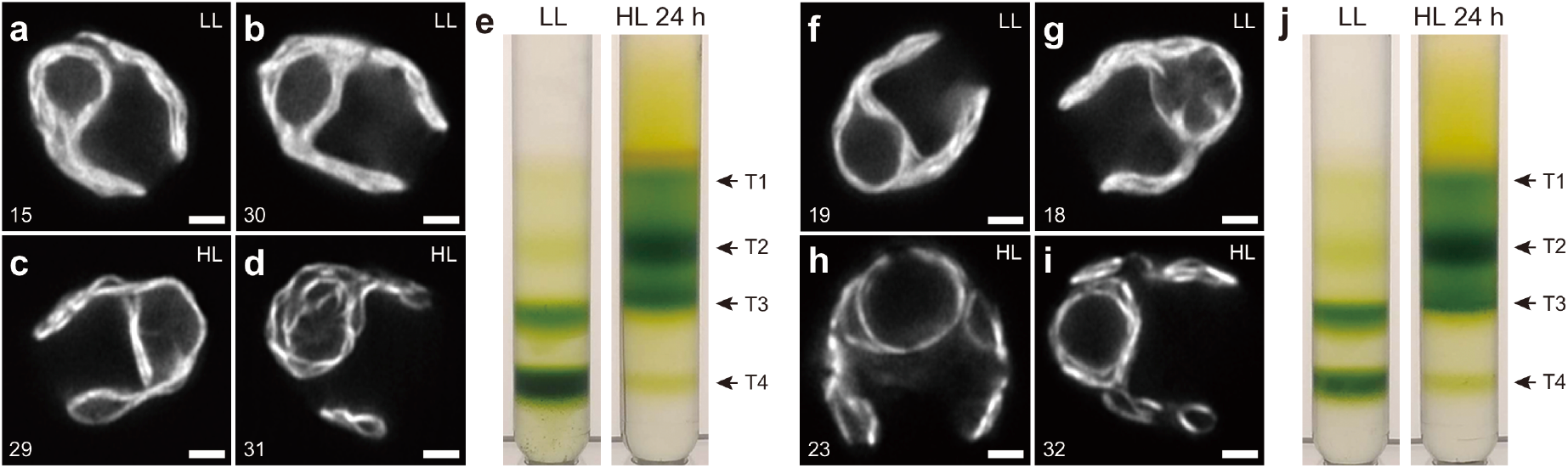
Thylakoid structural changes after HL acclimation in *npq4 lhcsr1* and *spa1-1*. Representative Airyscan images of the *npq4 lhcsr1* mutant cells acclimated to LL (**a, b**) and HL (**c, d**). **e.** Analytical sucrose density-dependent membrane fractionation using the *npq4 lhcsr1* mutant cells acclimated to LL and HL were gently disrupted (a total of 0.5 mg Chl) and separated by sucrose step-gradient centrifugation (see Methods for details). Representative Airyscan images of the *spa1-1* mutant cells acclimated to LL (**f, g**) and HL (**h, i**). **j.** Analytical sucrose density-dependent membrane fractionation using the *spa1-1* mutant cells acclimated to LL and HL were gently disrupted (a total of 0.5 mg Chl) and separated by sucrose step-gradient centrifugation (see Methods for details). Numbers (in a-d, f-i) indicate slice numbers of z-stack images. Scale bars = 2 μm. Different membrane fractions were indicated as T1-T4.

The results using *npq4 lhcsr1* suggest that thylakoid structural changes do not directly influence the qE capacity. To test this further, we used the *spa1-1* mutant, in which LHCSR accumulates at high levels even under LL (Gabilly *et al.*, 2019). While the stromal space around the pyrenoid was already enlarged before HL acclimation in *spa1-1* (Figure 5f, g), TEM images of LL-grown *spa1-1* cells revealed that the enlarged stromal space around the pyrenoid was likely a consequence of high starch accumulation in the pyrenoid, which was not observed in WT (Figure S1). After HL acclimation, similar structural changes were observed in *spa1-1* relative to WT (Figure 5h, i). Analytical density-dependent fractionation analysis still showed results similar to WT—more T4 fraction in LL and an increase in T1, T2, and T3 fractions after HL acclimation (Figure 5j). Taken together, these results reveal that LHCSRs in thylakoid membranes are not sufficient to cause the membrane unstacking observed during HL acclimation. The fact that LL-grown *spa1-1* has a high NPQ capacity (Gabilly *et al.*, 2019), albeit at reduced levels relative to HL-acclimated cells, also demonstrates that LHCSR-mediated induction of NPQ can at least partly occur in the absence of the thylakoid membrane structural rearrangement associated with HL acclimation. Therefore, LHCSRs are neither necessary nor sufficient to induce the thylakoid membrane structural rearrangement associated with HL acclimation, and LHCSR-mediated NPQ can operate, at least partly, in the absence of these structural rearrangements.

An alternative consideration to an LHCSR-dependent signal for thylakoid membrane structural rearrangements in HL is that they might relate to photosynthetic electron transport itself. To investigate whether the HL-induced thylakoid structural changes are dependent on photosynthetic electron transport, we treated *C. reinhardtii* cells with 3-(3,4-dichlorophenyl)-1,1-dimethylurea (DCMU), which blocks electron transfer from PSII to PQ, inhibiting photosynthetic linear electron transport. Interestingly, the results demonstrated that the accumulation of T1 and T2 fractions was suppressed by DCMU treatment, and accumulation of the T4 fraction increased relative to samples without DCMU (Figure 6a). These samples were harvested after 15 min of HL to limit the severe photodamage that HL induces in DCMU-treated cells. However, 15 min was sufficient to induce thylakoid structural changes similar to that observed after 24 h HL treatment in untreated cells (Fig. 4a-b).

PQ remains oxidized upon DCMU treatment. The PQ redox state is known to affect state transitions in *C. reinhardtii*, which maintains the energy balance between PSI and PSII by dynamically redistributing excitation energy absorption between the two photosystems via reorganization of LHCII (Minagawa, 2011, Rochaix, 2011, Goldschmidt-Clermont and Bassi, 2015). DCMU-treatment locks cells in the so-called “state 1 conditions” in which LHCII remains unphosphorylated (Allen *et al.*, 1981), and excitation energy is preferentially transferred to PSII (Finazzi *et al.*, 2002, Iwai *et al.*, 2008).

**Figure 6.**
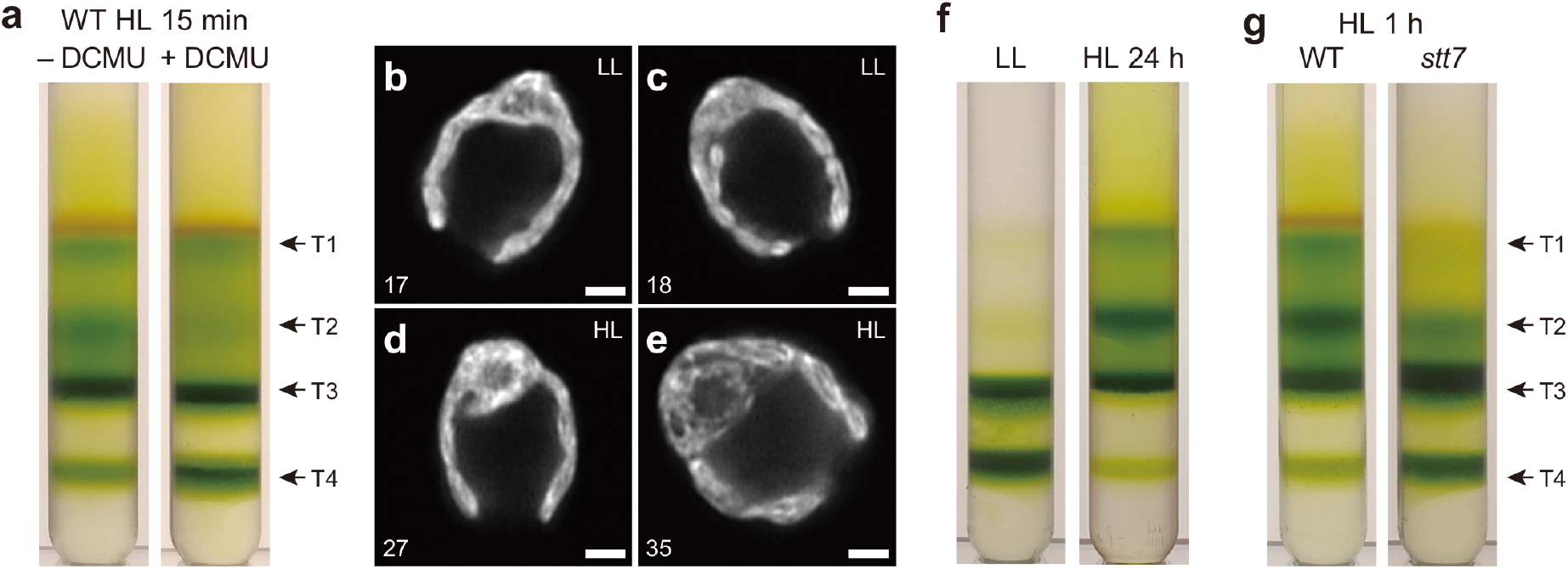
The effect of photosynthetic electron transfer and state transitions. **a.** Analytical sucrose density-dependent membrane fractionation using WT cells treated with DCMU during 15 min of HL treatment were gently disrupted (a total of 0.5 mg Chl) and separated by sucrose step-gradient centrifugation (see Methods for details). Different membrane fractions were indicated as T1-T4. **b.** Representative Airyscan images of the *stt7-9* mutant cells acclimated to LL (**b, c**) and HL (**d, e**). Numbers indicate slice numbers of z-stack images. Scale bars = 2 μm. **f.** Analytical sucrose density-dependent membrane fractionation using the *stt7-9* mutant cells acclimated to LL and HL (for 24 h) were gently disrupted (a total of 0.5 mg Chl) and separated by sucrose step-gradient centrifugation. **g.** Analytical sucrose density-dependent membrane fractionation using the WT and *stt7-9* mutant cells acclimated to HL for 1 h were gently disrupted (a total of 0.5 mg Chl) and separated by sucrose step-gradient centrifugation.

To test whether the HL-induced structural changes are related to state transitions, we used the *stt7-9* mutant, which lacks the Stt7 kinase that phosphorylates LHCII (Depege *et al.*, 2003) and thus does not perform state transitions. Intriguingly, there were less pronounced structural changes between LL- and HL-acclimated *stt7-9* cells than that observed in WT (Figure 6b-e). This result is in agreement with the observation in *Arabidopsis*, in which LHCII phosphorylation causes dynamic thylakoid membrane stacking (Fristedt *et al.*, 2009, Wood *et al.*, 2019). However, analytical density-dependent fractionation analysis still showed similar results to WT (Figure 6f). Because the state transition is a short-term light acclimation mechanism, HL treatment for 24 h might cause other long-term acclimation mechanisms. To observe the short-term effect on thylakoid structural changes, we treated both WT and the *stt7-9* mutant under HL for 1 h and analyzed them by density-dependent fractionation analysis. Interestingly, the result showed that the *stt7-9* cells had much less T1 and T2 and more T3 and T4 than in the WT (Figure 6g). Thus, at least for a short-term response to HL, LHCII phosphorylation is necessary for the membrane unstacking as observed in WT. These results indicate that Airyscan imaging and analytical density-dependent fractionation actually characterize aspects of thylakoid membrane structures that are at different spatial levels—the former visualizes overall membrane structural shapes, while the latter specifically distinguishes the level of membrane stacking.

## DISCUSSION

Our observations demonstrated that macroscale structural changes of thylakoid membranes occur in *C. reinhardtii* during HL acclimation (Figures 1-4). According to the results using *npq4 lhcsr1* and *spa1-1*, the HL-induced structural changes are not directly induced by the accumulation of LHCSR proteins in thylakoid membranes (Figure 5). These results also reconfirmed that high NPQ capacity in *C. reinhardtii* is strongly correlated with the LHCSR protein level in thylakoid membranes but not with the macroscale structural changes of thylakoid membranes. Notably, the NPQ capacity of *spa1-1*, which accumulates high LHCSR proteins even under LL, was higher in LL but still increased further after HL acclimation (Gabilly *et al.*, 2019). Thus, while LHCSR is neither necessary nor sufficient to induce thylakoid structural rearrangements, the high NPQ capacity of HL-acclimated cells may well depend on the thylakoid structural changes during HL acclimation.

DCMU not only blocks linear electron transport but also the HL-induced expression of *LHCSRs* (Petroutsos *et al.*, 2011, Maruyama *et al.*, 2014). Interestingly, the DCMU-treated cells did not show the structural changes under HL, at least during a short period of time of HL exposure (Figure 6a). The reduced state of the PQ pool also induces LHCII phosphorylation (Finazzi, 2005, Rochaix, 2007). Although the lack of LHCII phosphorylation in state transitions does not directly affect NPQ capacity (Girolomoni *et al.*, 2019), it has been shown that LHCSR can move to PSI during state transitions (Allorent *et al.*, 2013). We observed that the lack of LHCII phosphorylation indeed prevented membrane unstacking in the beginning of HL treatment (Figure 6g). Thus, it seems likely that thylakoid membrane protein reorganization in a short-term HL acclimation depends on LHCII phosphorylation. However, prolonged HL conditions (e.g. 24 h) eventually generate a similar level of unstacked membranes to that of WT in the *stt7-9* mutant (Figure 6f). Unexpectedly, Airyscan images did not show the pronounced HL phenotypes in the *stt7-9* mutant after 24 h of HL treatment (Figure 6d, e), suggesting that the macroscale membrane architecture and the level of membrane stacking need to be determined separately.

Another possible relationship worth considering is the thylakoid membrane structures and how they facilitate and influence protein assembly and insertion into thylakoid membranes. During HL acclimation, the newly translated LHCSR proteins need to be inserted in a way that excitation energy is efficiently transferred from LHCII to be dissipated by LHCSR. The base region of the *C. reinhardtii* chloroplast near the pyrenoid is known as the translation zone, in which protein translation and assembly as well as Chl biosynthesis occur (Schottkowski *et al.*, 2012, Sun *et al.*, 2019). Our imaging results showed thinning of the overall thylakoid structures, including the translation zone, and the enlarged stromal space at the base region during HL acclimation (Figs. 2, 3). To be functional, newly synthesized LHCSR proteins need to be assembled with Chls and carotenoids (Bonente *et al.*, 2011). The HL-induced structural changes of thylakoid membranes might be coordinated with protein assembly and insertion mechanisms, and the membrane unstacking may enable newly synthesized photoprotective proteins such as LHCSR to have access to membranes to be inserted. This would provide a mechanistic explanation for why ectopic expression of LHCSR in LL is not sufficient to induce levels of NPQ equal to HL-grown cells (Gabilly *et al.*, 2019).

It is worth mentioning the limitations of imaging analysis by observing Chl fluorescence. It is practical to assume that the structures based on Chl fluorescence imaging reflect thylakoid membranes because the membranes are filled with Chl pigments. However, Chl fluorescence emission from PSI is difficult to observe at room temperature, especially with confocal microscopes, due to its higher excitation trapping than PSII (Trissl and Wilhelm, 1993). Also, fluorescence intensity frequently fluctuates due to several factors, including NPQ and photosynthetic electron transport, that can influence the interpretation of observed structural differences in thylakoid membranes due to different intensities of Chl fluorescence between LL- and HL-acclimated cells. Therefore, it is necessary to use TEM to confirm the structural changes in thylakoid membranes observed by Airyscan microscopy (Figure 3). The TEM images also reveal the similar structural differences in thylakoid membranes and indicate that Airyscan microscopy is capable of visualizing the overall thylakoid structures sufficiently to reveal the differences between the LL- and HL-acclimated cells. It should be noted that, although large stromal spaces between membranes are more visible in the HL-acclimated cells than that of the LL-acclimated cells, it also appears that a lot of membranes in the HL-acclimated cells are associated closely with each other (Figure 3e, g, h). This might suggest a possible alteration of refractive indices of the membrane, which would affect how much light is absorbed by LHC proteins (Capretti *et al.*, 2019). However, because of the chemical fixation and negative staining done in conventional TEM, it is difficult to determine whether changes in such membrane compartmental spaces are correlated with native conditions. Although bioimaging techniques provide images at lower resolution than do EM techniques, it is feasible to evaluate a large number of samples, which is inherently practical to determine statistically supported characteristics when examined under EM. Therefore, our observation will be helpful when microscopic details of the thylakoid compartment, stacking, and protein organization will be examined by advanced cryo-EM techniques.

## EXPERIMENTAL PROCEDURES

### Strains, growth conditions, and HL treatment

*C. reinhardtii* WT strain 4A+ (mt+, 137c background; CC-4051), *npq4 lhcsr1*, and *spa1-1*, and *stt7-9* mutants were grown in Tris-acetate-phosphate liquid media (Harris *et al.*, 1989) as described previously (Niyogi *et al.*, 1997). The culture was adjusted to ~1 μg Chl/mL and incubated on a shaker under ~30 μmol photons m^−2^ s^−1^ (LL) at 25°C for 3 d. For HL treatment, the LL-acclimated cells were diluted to 1 μg Chl/mL (for 24 h treatment) or 2 μg Chl/mL (for 15 min and 1 h treatments) in fresh TAP liquid media and incubated under ~350 μmol photons m^−2^ s^−1^ at 25°C for specified duration of time. For the 24-h incubation under HL, the culture was diluted again after 6 h of HL incubation to maintain a low cell density. Chl concentration was measured as described previously (Porra *et al.*, 1989).

### Airyscan microscopy

*C. reinhardtii* cells were observed using a Zeiss LSM 880 microscope equipped with the Airyscan detector with a Zeiss Plan-Apochromat 63×/1.4 NA DIC M27 Oil objective. Chls were excited with 633 nm laser, and fluorescence was acquired through a 645 nm longpass filter. Image acquisition and analysis were done under the full control of ZEN software (Zeiss) and ImageJ software (US National Institutes of Health, https://www.nih.gov/).

### TEM

*C. reinhardtii* cells acclimated under LL and HL 24 h were centrifuged, resuspended, and fixed with 2% glutaraldehyde in TAP liquid media for 24 h at 4 °C in the dark. The fixed cells were washed and post-fixed with a sodium cacodylate buffer containing 1% (w/v) osmium tetroxide and 0.8% (w/v) potassium ferricyanide for 2 h. The fixed cells were rinsed with cacodylate buffer for 10 min twice. The fixed cells were dehydrated with increasing concentrations of acetone (35-100%). The dehydrated cells were then infiltrated and embedded in resin. Sections were cut to approximately 500 μm in diameter and 60 nm in thickness. The thin sections were collected on Maxtaform copper slot grids (2 × 1-mm oval hole) that had been coated with 0.5% formvar. Sections were dried and post-stained with 2% uranyl acetate for 7 min, followed by lead citrate for 7 min. The sections were dried and examined using a JEOL 1200 EX transmission electron microscope at 100 kV.

### AFM

Height of isolated thylakoid membrane was measured as described previously (Iwai *et al.*, 2013). Briefly, a freshly cleaved mica surface was treated with adsorption buffer (10 mM Tris-HCl (pH 7.3), 150 mM KCl, 20 mM MgCl2). Then, isolated thylakoid membranes (0.25 μg Chl/μL) were added to the adsorption buffer on the mica surface and incubated for 10 min. After gentle wash with MilliQ water, the membranes were observed by using a commercial MFP-3D stand-alone AFM in tapping mode in air (Asylum Research). Silicon cantilevers with a length of 240 μm (*k* = 2 N/m; OMCL-AC240TS-C2, Olympus) were used. Averaged height was calculated from the membrane areas of at least 50 μm^2^.

### Analytical sucrose density-dependent membrane fractionation

*C. reinhardtii* cells were collected at 1,870 × *g* and 4°C for 5 min (JLA9.1000, Beckman Coulter). The cells were resuspended at 0.125 mg Chl/mL with disruption buffer containing 25 mM MES-NaOH (pH 6.5), 0.33 M sucrose, 1.5 mM NaCl, 0.2 mM benzamidine, and 1 mM ε-aminocaproic acid at 4°C. The cells were disrupted at 5 kpsi once using the MC Cell Disruptor (Constant Systems Ltd, Northants, UK). The disrupted cells (0.5 mg Chl in 4 mL) were loaded onto sucrose step-wise gradient, containing 0.9, 1.2, 1.5, and 1.8 M sucrose with 25 mM MES-NaOH (pH 6.5) and 1.5 mM NaCl (2 mL/each sucrose layer). The thylakoid membranes with different densities were analytically separated at 125,000 × *g* and 4°C for 1 h (SW 41 Ti, Beckman Coulter).

### Fluorescence emission spectroscopy at 77 K

The sample obtained by analytical sucrose density-dependent membrane fractionation was placed in a glass tube and frozen in liquid nitrogen. Fluorescence emission was recorded at 77 K using FluoroMax-4 spectrophotometer (Horiba Scientific). Excitation wavelength was 440 nm with a 2-nm slit size. Emission wavelength measured was from 650 to 800 nm with a 2-nm slit size. Fluorescence emission for each sample was recorded consecutively three times to obtain averaged spectra.

## ACKNOWLEDGEMENTS

We thank Christopher R. Baker, Setsuko Wakao, and Valle Ojeda for their critical reading of the manuscript; Holly Aaron and Feather Ives at the Molecular Imaging Center at University of California, Berkeley for the technical setup for Airyscan microscopy; and Reena Zalpuri at the Electron Microscope Laboratory at University of California, Berkeley for assistance in EM sample preparation and data collection. This work was supported by the U.S. Department of Energy, Office of Science, through the Photosynthetic Systems program in the Office of Basic Energy Sciences. K.K.N. is an investigator of the Howard Hughes Medical Institute.

## CONFLICT OF INTEREST

The authors declare no conflict of interest.

## SUPPORTING INFORMATION

**Figure S1.**
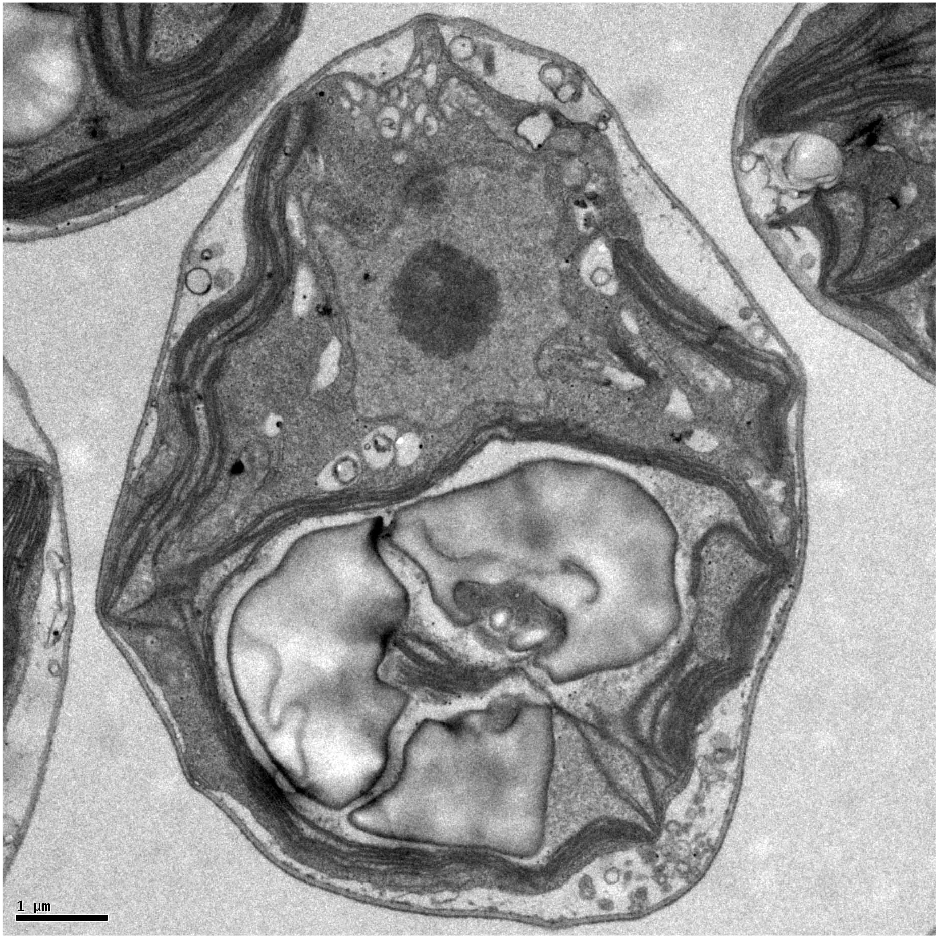
The representative TEM image of the *C. reinhardtii spa1-1* cell acclimated to LL conditions. Scale bar = 1 μm.

## Notes

### Competing Interest Statement

The authors have declared no competing interest.

